# Championing Health Promotion through Group Patient and Family Education Activities in the Outpatient Department of a Tertiary Care Hospital

**DOI:** 10.1101/813360

**Authors:** Mahmoud Ibrahim Almahameed, Shadi F. Kakish, Amani Abu-Shaheen

## Abstract

**Background:** Health education involves not only providing relevant information but also facilitating health-related behavior change. Patient education during the waiting period plays an essential role in promoting health and improving patient satisfaction rates.

**Objective:** To promote health by educating patients and their companions during their waiting period at the King Fahad Medical City (KFMC) Hospital, Riyadh, with the help of nurses and assess their satisfaction rate.

**Methods:** A cross-sectional study, including patients and their companions in the waiting rooms of the various OPD’s, was conducted. Group teaching sessions were conducted, followed by the assessment of patient and nurse satisfaction rate, and compliance rate using evaluation forms. Pre-implementation surveys were conducted to assess the drawbacks, based on which procedure was modified, and feedback was recorded using an electronic iPad. Data were collected with the help of questionnaires filled at the end of each teaching session and analyzed using descriptive analysis. Results of both the assessment and re-assessment were compared in terms of the number of attendees.

**Results:** There was an improvement in number of attendees (4362 vs. 3392), patient satisfaction rate (90% vs. 78%), nurse satisfaction rate (83.09% vs. 33.18%) and compliance rate (100% vs. 65%) post-modifications. Few loopholes in terms of inappropriate environment and time of the lecture were also observed.

**Conclusion:** The health educators can assist people in achieving their health goals by organizing structured health education activities, providing appropriate health education materials, and effective training. However, the satisfaction rates of people and health educators should be assessed by conducting routine feedback surveys.

## Introduction

According to the World Health Organization, "Health promotion is the process of enabling people to increase control over and to improve their health" ^1^. It motivates individuals to take initiatives in health literacy and multisectoral intervention to improve healthy habits ^2^.

Hospitals play a critical part in health promotion, advancing wellbeing, anticipating illness, and providing rehabilitation services ^3^. Nurses and clinical health educators are in the best place to meet patients and their families’ health promotion requirements ^4^. Responsibility for health promotion in health facilities is equally distributed between the people, community organizations, healthcare professionals, health organizations, and government ^5^.

The patient education approach is an efficient way to improve general patient satisfaction and ultimately enhance general demographic health status ^6^. As per the Joint Commission International (JCI) Accreditation Standards for Hospitals, the hospital should provide patient education to help involve patients and their families in care decisions and care procedures ^7^. However, educational methods used to impart knowledge should consider the patient’s and their family’s values and preferences to allow enough interaction among the patient, family, and staff ^7^.

Patient education during the waiting period plays an essential role in meeting patients’ learning requirements and can influence patient satisfaction rates ^8^. Also, assessing patient satisfaction is crucial to health educators, doctors, hospital administrators, and patients themselves to guarantee that healthcare requirements are met and preserved ^6^.

According to the General Authority for Statistics, Kingdom of Saudi Arabia, the population of Saudi Arabia was more than 33 million in the year 2018. With the rising trend in population, the burden on healthcare has also increased. The government of Saudi Arabia has made tremendous efforts to improve healthcare through health education ^9^. Under King Fahad Medical City (KFMC) Strategic Plans 2015-2020, the KFMC in Riyadh, adopted some key strategies to provide excellence in health management and patient’s experience and utilized resources in initiating group teaching activities in the outpatient department (OPD) ^10^. Thus, the present study aims to promote health by educating patients and their attendees during their waiting period in OPD at KFMC hospital with the help of healthcare professionals.

## Materials and methods

### Study design

A cross-sectional study was performed at KFMC Hospital in Riyadh, Saudi Arabia between, July 2017 and December 2018 to enhance health education among patients and their companions.

### Study population

The participants included in the study consisted of adult patients with booked appointments and their companions present in the waiting areas of various OPD’s of the hospital.

### Recruitment and sample size estimate

The study participants were randomly approached and invited to take part in this study from the waiting area of the OPD.

According to the KFMC’s target through a focus group method, the study aimed to target 2% of the total patients and their companions who visited the OPD per month.

### Study procedure

The study procedure consisted of four stages, *i.e.*, group teaching, assessment, intervention/ modification, and re-assessment.

#### Group teaching

The healthcare professionals of the hospital prepared a list of educational/ health-related topics to be discussed with the patients and their companions present in the waiting area of OPD’s. The group teaching sessions involved discussions on those health-related topics by nurses. On completion of discussion, evaluation sheets consisting of questions were distributed to the patients and their companions present in the waiting area of the OPDs. Once the forms were filled, they were collected from them for further analysis.

#### Questionnaire / Assessments

Questionnaires were used to evaluate the efficacy of patient and family education (PFE) activities, and the assessment was made for the following parameters:

- Number of participants: The number of patients and their companions participating in the teaching sessions were recorded monthly. The data was summed-up to determine the total number of participants.
- Patient experience: A questionnaire consisting of 14 questions was prepared to evaluate the patient experience. All the questions, except Q.9, included two options, Yes or No.
- Patient and family satisfaction: A questionnaire consisting of four questions and four options (not satisfied, satisfied, totally not satisfied, and very satisfied) was prepared.
- Staff satisfaction: A questionnaire consisting of 11 questions was prepared. All the questions except Q.1 included two options, Yes or No.
- Compliance with planned educational topics: Data was collected from the questionnaires to determine the compliance of planned educational topics for lectures.

#### Intervention/ modification

Pre-implementation surveys: Pre-implementation surveys are conducted to achieve an enhanced understanding of the potentials and weaknesses of the study and guide project development ^23, 24^. The pre-implementation surveys were conducted by involving 20 staff nurses/ health care assistants and 104 patients to identify the factors, if any, responsible for the low satisfaction rate observed after assessment of patients’ and staff experience.

The FOCUS-PDCA quality model was selected to analyze and improve the drawbacks reported during the previous assessment. The term FOCUS-PDCA stands for the words to find, organize, clarify, understand, select, plan, do, act, and check results. The FOCUS-PDCA model is a useful performance improvement strategy developed by the Hospital Corporation of America. It has been applied successfully in various domains such as nursing, surgery, laboratories, trauma, and used by many healthcare organizations ^12^.

Identification of loopholes: Root cause analysis and rigorous brainstorming were performed on the pre-implementation survey results to identify the drawbacks/ challenges. Root cause analysis is a method for analyzing adverse events. It is widely used as an error analysis tool in healthcare ^11^.

Modifications: To improve the health educator skills, participants’ experience, and their satisfaction rates, the following measures were taken:

- Two workshops were conducted for Arabic health educators to increase their competence in the delivery of PFE sessions.
- New health education materials were prepared by the OPD nursing team
- Approved new health education materials were uploaded on the KFMC iPad for the accessibility of resources to all staff and participants
- Electronic surveys were also uploaded on iPad to expedite and receive prompt feedback from patients in response to the group teaching sessions
- Structured group teaching activities were organized
- The efficacy of the group teaching activities was monitored through surveys
- Monthly meetings were conducted and attended by the multidisciplinary team

Besides, the procedure of group teaching was also modified.

The topics with unavailable material were not modified/ replaced with the other. However, approved educational material related to that topic was downloaded from the hospital’s intranet. The session was only conducted if the number of participants was more than four or about 4-10. The staff introduced themselves, the topic, and the duration of the session to the participants before conducting the session. On completion, the staff instructed the participants on how to fill the feedback forms through the iPad.

#### Re-assessment

The patients/ staff experience and satisfaction were determined using the same questionnaires but were filled electronically using the hospital’s iPad. The results obtained from both the surveys were compared to assess change/ improvement in the experience, satisfaction, and compliance. Therefore, the study endpoints were to determine the increase in the number of patients and their companions, improvement in patient experience, patient and family satisfaction rate, staff and nurse satisfaction rate, and compliance rate to planned educational topics after modifying the group teaching procedure.

### Data analysis

Data were analyzed descriptively and were presented as number and percentages using SPSS version 21.0 statistical software (IBM Corp., Armonk, NY, USA).

## Results

### Study population

Overall, a total of 7754 patients and their companions participated in health education activities.

### Assessment

#### Number of patients and their companions receiving health education

The number of patients and their companions who received education were 3,392. The focus group target was achieved, averaging 2.69% of the total patients and their companions who visited OPD per month.

#### Patient experience of health education activities

Overall, 63.86% of patients reported positive patient experience to the health education activities conducted. The participants believing that educational sessions influentially affect them, and the society was 82.18% (n=2788). A total of 76.24% (n=2586) patients stated that health practitioners could answer their queries. Also, 86.14% (n=2922) of the participants shared the information received amongst their families and community (Table 1).

**Table 1:** Questionnaire for Evaluating Patient Experience.

| Sr. No | Questions | Satisfaction Rate*<br>(%) |  |
| --- | --- | --- | --- |
|  |  | Assessment | Re-assessment |
| 1. | Do you know that there are educational lectures by nurses while coming to the OPD? | 1478 (43.56) | 1552 (35.58) |
| 2. | Have you been invited before to attend an educational lecture on visiting the OPD? | 1343 (39.60) | 1552 (35.58) |
| 3. | Is there any announcement of the educational campaign? | 1310 (38.61) | 1804 (41.35) |
| 4. | Do you think that the educational sessions will have an influential effect on the patient and society? | 2788 (82.18) | 3817 (87.50) |
| 5. | Can the health practitioner (nurse) answer your questions? | 2586 (76.24) | 3859 (88.46) |
| 6. | Do the nurses give you useful teaching materials? | 2150 (63.37) | 3020 (69.23) |
| 7. | Can you learn and gain new information when you visit the clinic? | 2687 (79.21) | 3565 (81.73) |
| 8. | Do you share the information with your family and community? | 2922 (86.14) | 4027 (92.31) |
| 9. | Are you satisfied with the current method for health education? | 2217 (65.35) | 2726 (62.50) |
| 10. | Do you get health information through social media like WhatsApp, Twitter, YouTube, Snapchat, Instagram? | 2653 (78.22) | 3146 (72.12) |
| 11. | Do you get health information through television? | 2317 (68.32) | 2852 (65.38) |
| 12. | Do you get health information during your visit to the OPD or Hospital? | 2452 (72.28) | 3565 (81.73) |
| 13. | You get health information through from the newspaper? | 1545 (45.54) | 1971 (45.19) |
| 14. | Do you get health information through relatives and other patients? | 1881 (55.45) | 3607 (82.69) |
| **Overall** |  | **2166 (63.86)** | **2933 (67.24)** |
| OPD: Outpatient Department; *: Number of patients giving favorable response. |  |  |  |

#### Patient and family satisfaction from health education activities

The patient and their companions reported a 78% satisfaction rate on the conduct of PFE activities. About 78.21% (n=2653) participants reported getting an answer to their queries. Approximately, 78.30% (n=2656) participants reported being very satisfied with the educational sessions in terms of duration. In terms of clarity of the educational sessions, 78.18% (n=2652) reported being very satisfied. The low satisfaction rates attributed to the unclear objectives, unanswered queries, and improper duration of the lectures (Table 2).

**Table 2:** Questionnaire for Evaluating Patient and Family Satisfaction

| Sr. No | Questions | Not satisfied |  | Satisfied |  | Totally not satisfied |  | Very Satisfied |  | Total |  |
| --- | --- | --- | --- | --- | --- | --- | --- | --- | --- | --- | --- |
|  |  | A<br>n (%) | RA<br>n (%) | A<br>n (%) | RA<br>n (%) | A<br>n (%) | RA<br>n (%) | A<br>n (%) | RA<br>n (%) | A<br>n (%) | RA<br>n (%) |
| 1 | Did we answer your questions? | 398<br>(11.73) | 2<br>(0.04) | 346<br>(10.20) | 454<br>(10.40) | 2<br>(0.05) | 6<br>(0.13) | 2653<br>(78.21) | 3900<br>(89.40) | 3392 | 4362 |
| 2 | Was it too long for you? | 398<br>(11.73) | 1<br>(0.02) | 352<br>(10.37) | 304<br>(6.96) | 1<br>(0.02) | 0 (0) | 2656<br>(78.30) | 4057<br>(93.00) | 3392 | 4362 |
| 3 | Were our message and goal clear to you? | 377<br>(11.11) | 7<br>(0.16) | 350<br>(10.31) | 333<br>(7.63) | 7<br>(0.20) | 5<br>(0.11) | 2652<br>(78.18) | 4017<br>(92.09) | 3392 | 4362 |
| 4 | Would you recommend us to others? | 378<br>(11.14) | 3<br>(0.06) | 347<br>(10.22) | 336<br>(7.70) | 3<br>(0.08) | 1<br>(0.02) | 2646<br>(78.00) | 4022<br>(92.20) | 3392 | 4362 |
| **Total** |  | **388** | **12** | **349** | **628** | **3** | **12** | **2652** | **4218** | **3392** | **4362** |
| A: Assessment, n: Number of Participants, RA: Re-assessment. |  |  |  |  |  |  |  |  |  |  |  |

#### Staff and nurse satisfaction from health education activities

The assessment reported an average staff-satisfaction rate of 33.18%. The factors determining the satisfaction rate were staff experience and educational barriers. One-fourth (25%, n=6) of the staff was satisfied with the process of PFE followed. About 35% (n=8) of the staff observed difficulty in getting the educational materials. Around 30% (n=7) of the staff reported that the patients responded to their call for the educational lectures. Also, 10% (n=2) of the staff reported difficulty in answering the patient’s queries. The major barriers observed by the staff included the lecture timings (60%), availability of educational materials (55%), venue (40%), gender preferences of the participants (35%), presentation skills (30%), and language (10%) (Table 3).

**Table 3:** Questionnaire for Evaluating Nurse Satisfaction

| Sr. No | Questions | Satisfaction Rate* |  |
| --- | --- | --- | --- |
|  |  | % |  |
|  |  | Assessment | Re-assessment |
| 1. | Are you satisfied with the current method of patient and family education? | 6 (25) | 19 (86) |
| 2. | Is it easy to get educational materials for the patients? | 8 (35) | 19 (86) |
| 3. | Are you able to answer the patient's questions and queries? | 2 (10) | 19 (86) |
| 4. | Do the patient's respond to your call for the educational lecture? | 8 (35) | 18 (81) |
| 5. | Do you distribute the educational materials before and after each lecture? | 7 (30) | 19 (86) |
| 6. | Is language a barrier? | 2 (10) | 21 (95) |
| 7. | Is the place of lecture a barrier? | 9 (40) | 13 (57) |
| 8. | Is the time of lecture a barrier? | 13 (60) | 17 (76) |
| 9. | Is the availability of educational materials a barrier? | 12 (55) | 20 (90) |
| 10. | Is the gender of the patient a barrier? | 8 (35) | 18 (81) |
| 11. | Is your ability to explain and presentation skills a barrier? | 7 (30) | 20 (90) |
| **Overall** |  | **7 (33.18)** | **18 (83.09)** |
| *: Number of nurses giving favorable response. |  |  |  |

#### Compliance to planned educational topics

The average compliance rate of the planned educational activities observed was 61.25%.

### Loopholes

The pre-implementation surveys conducted after the assessment highlighted the shortcomings prevalent during the group teaching session. The significant shortcomings included viz.,

- Unavailability of patient and family education materials
- Low staff satisfaction with regards to the current process of PFE
- Disturbances during the PFE lectures (Queuing system, noise, etc.)
- The inability of staff to answer specific patient queries
- The improper venue of the PFE lecture (Overcrowded waiting areas)
- Unclear time and schedule for the PFE lecture
- Patient gender
- Insufficient training and poor presentation skills for the staff nurse
- The limited number of Arabic language speakers.

The drawbacks are presented in detail as a fishbone diagram (Figure 1) and Pareto chart (Figure 2).

**Figure 1:**
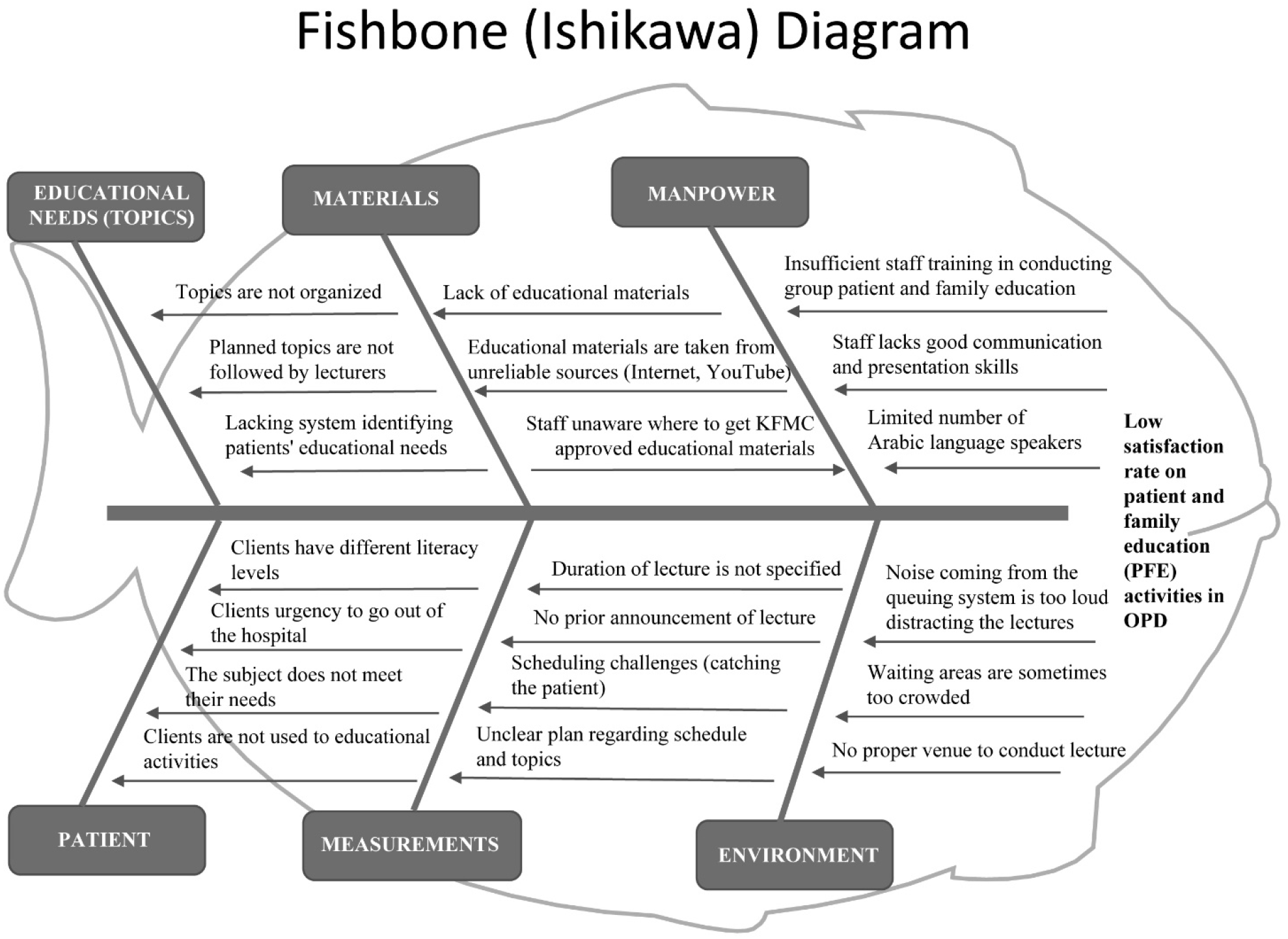
Fishbone diagram

**Figure 2:**
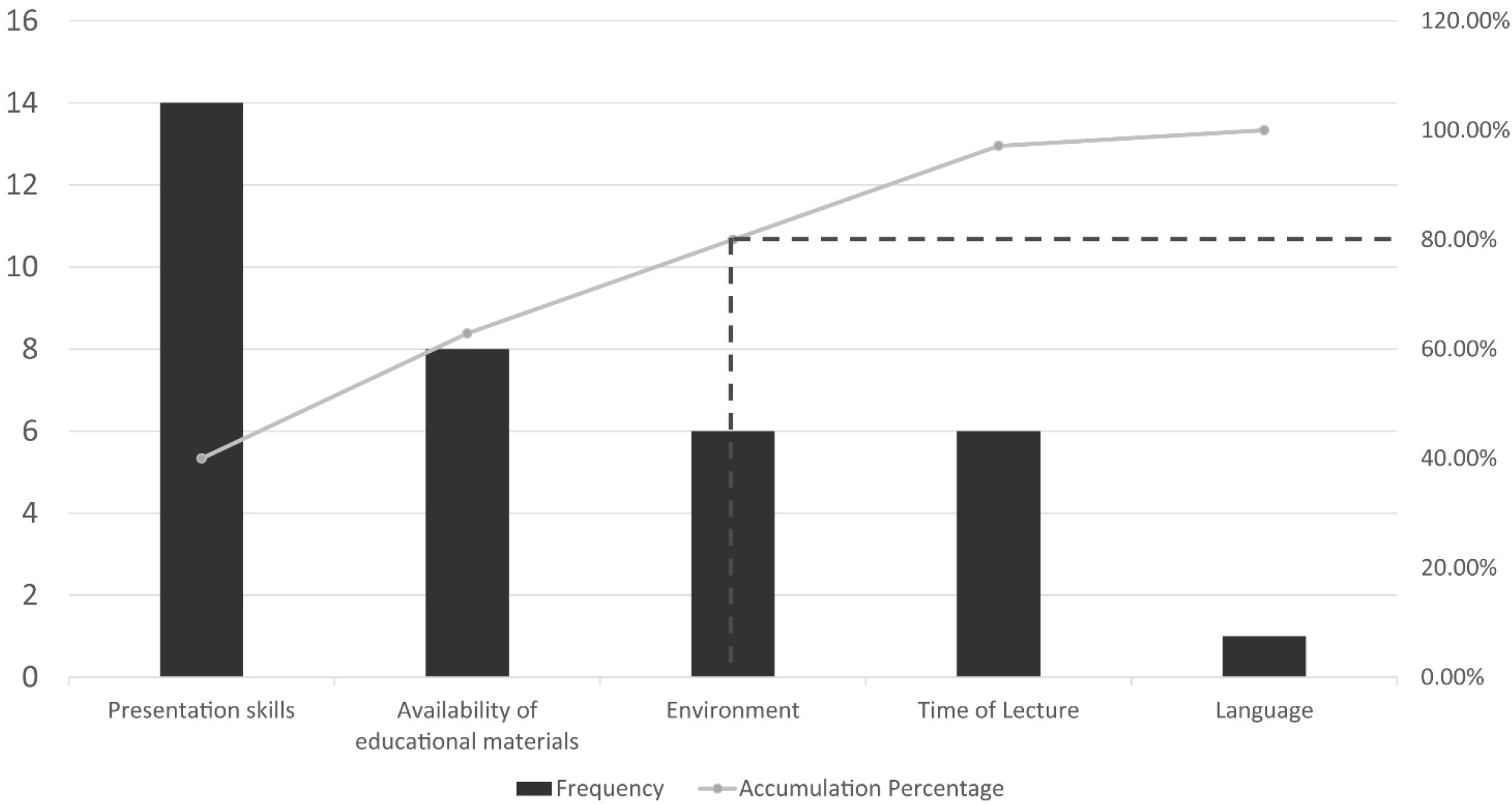
Pareto Chart on Low Satisfaction Rating (Assessment).

### Re-assessment

After modifying the group teaching procedure and adapting the interventions, the following improvements were observed:

#### Increase in the number of patients and their companions

The number of patients and their companions who received education increased to 4,362. Though the focus group target was achieved in both the assessments, there was an increase in the number of participants, averaging 3.07% of the total patients and companions.

#### Patient experience

The re-assessment revealed an improvement in the patient experience (67.24%) with the modified health education activities. The participants believing the educational sessions influence the patient and society increased to 87.50% (n=3817) post-modification. The percentage of health practitioners answering participants’ queries also increased to 88.46% (n=3859) About, 92.31% (n=4027) of the participants shared the information received amongst their families and community. Table 1 compares the results of both the assessments for patient experience.

#### Patient and family satisfaction from health education activities

An increase in the patient and family satisfaction rate (90%) was noted on re-assessment (Figure 3). Approximately 3900 (89.40%) participants reported getting answers to their queries, while 4057 (93%) participants reported being very satisfied with the educational sessions in terms of the duration of the sessions. Overall, the participants were more satisfied with the health education activities in the OPD after modifications (Table 2).

**Figure 3:**
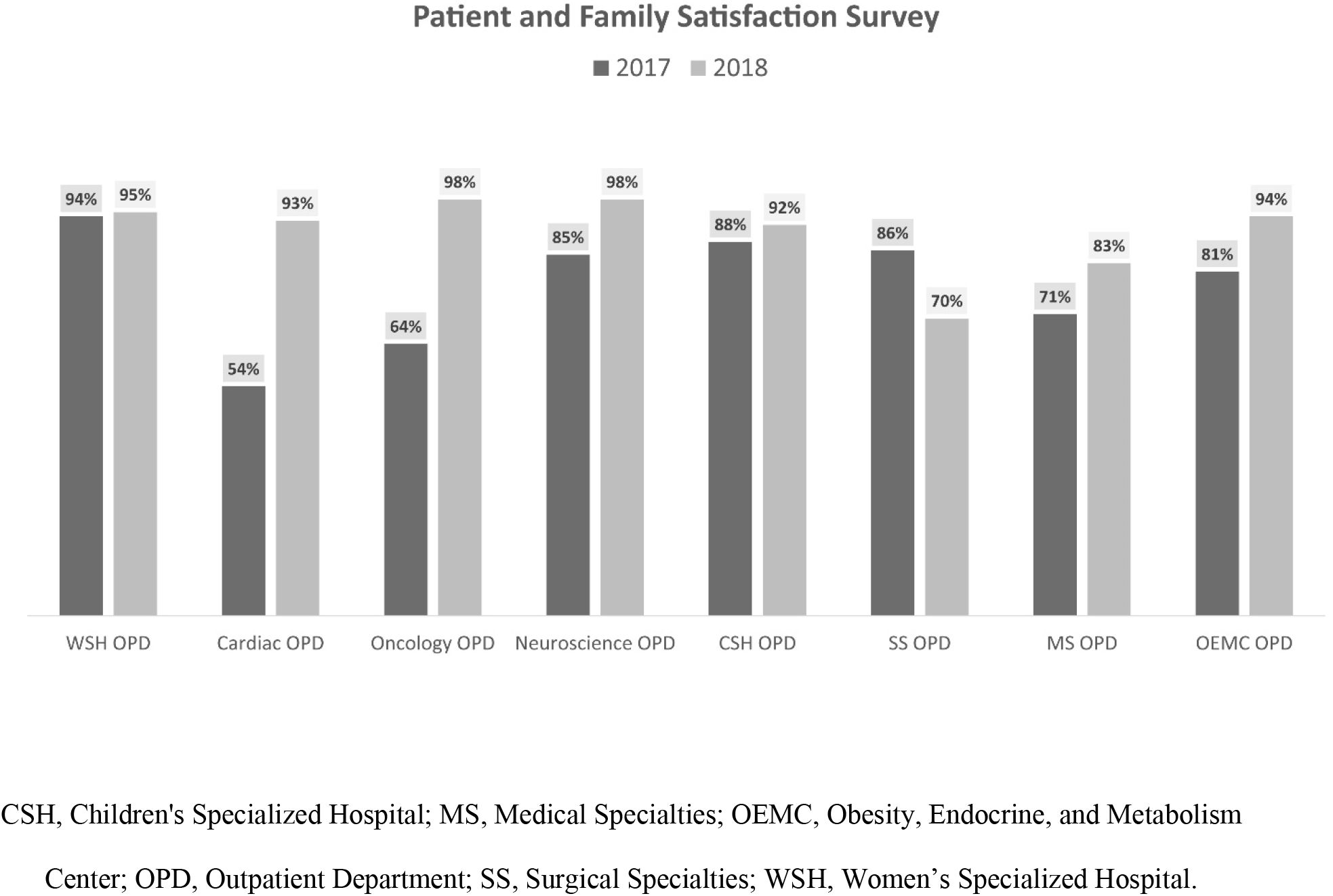
Patient and Family Satisfaction Survey Rate.

#### Staff and nurse satisfaction from health education activities

With the improvisation in the PFE methods, the staff experience improved significantly, with 86% satisfaction rate. In terms of answering the participant’s queries, 86% (n=19) of the staff members were satisfied. About 81% (n=18) of the staff had positive feedback concerning the participant’s attention, and 86% (n=19) reported on the ease in availability of educational materials and being able to answer patient queries. Besides, the improved process and modifications implemented after the pre-implementation surveys also helped overcome the staff barriers related to lecture timings, availability of educational materials, venue, gender, presentation skills, and language previously encountered (Table 3). The overall staff satisfaction rate post-modifications increased to 83.09% (n=18).

#### Compliance to planned educational topics

With the availability of patient education materials, the compliance to planned educational topics in the OPD increased by 38.75%, thereby strengthening the project structure.

### Loopholes

Despite the modifications made and the increase in satisfaction rate achieved on re-assessment, there were few loopholes observed such as inappropriate environment and prolonged or short duration of PFE lecture. The challenges were identified with the help of pareto charts based on the Pareto principle (Figure 4).

**Figure 4:**
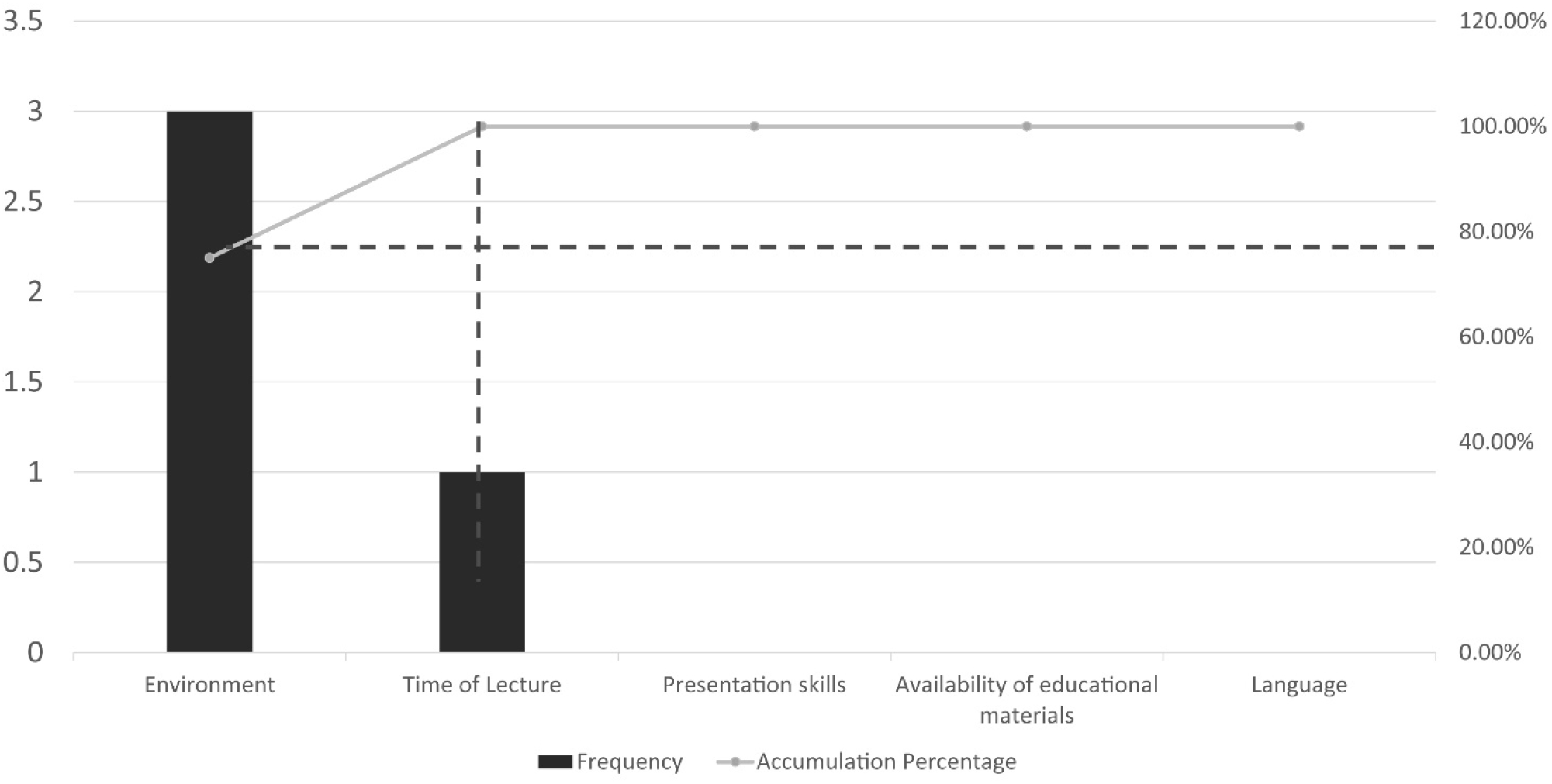
Pareto Chart on Low Satisfaction Rating (Re-assessment).

## Discussion

Education is a systematic, sequential, logical, scheduled course of action comprising of both teaching and learning ^13^. In terms of health, education involves providing relevant health-related information, influencing individual health choices, facilitating health-related behavior change, and increasing health awareness ^7, 14^. Health education, when provided to patients, demonstrate improved health outcomes and quality of life ^15^. The key role players involved in providing health education to patients and their families are health care professionals ^14, 16^. Educational activities in a hospital setting not only positively affect the patients but also benefit their families, providers, and society ^17^. Numerous studies have also highlighted the benefits of patient education activities in a hospital setting ^18–22^.

The KFMC hospital adopted some key strategies per the KFMC Strategic Plans 2015-2020, to initiate a group-teaching project in the OPD. Keeping in mind the low satisfaction rates of patients and healthcare professionals from the PFE lectures, pre-implementation surveys were conducted to assess the loopholes. Literature also reports the use of pre-implementation surveys to achieve an enhanced understanding of the potentials and weaknesses of the study and guide project development ^23, 24^. In the present study, pre-implementation surveys reported unorganized topics, schedules, unavailability of educational materials, communication gaps, lack of presentation skills, and improper venue as the significant shortcomings. Therefore, suitable modifications were made in the health education activities to improve the previously low satisfaction rates observed. These include implementing electronic surveys, producing additional new health education materials, staff training, organizing structured group education sessions, etc.

Group patient education activities have various benefits over individualized patient education methods in terms of cost-effectiveness, lesser workload, patients’ preference for discussing topics during group sessions, and reduction in repetition from individualized sessions ^19–21^. Asiri et al., 2013 conducted a study to evaluate patient satisfaction with various health educational services (individualized face-to-face sessions, group teaching, education exhibition, advice given in the waiting area, receiving printed materials) provided in different primary health care centers in Riyadh. The study reported the group teaching method as the most satisfactory method of patient education with a satisfaction rate of 87.2%. The results of our study are consistent with the results observed by Asiri et al., 2013 reporting a satisfaction rate of 78% (pre-modification) and 90% (post-modification) with the group health education activities ^6^. Various other studies conducted by Wilson, 1997, Trento et al., 2001, Merekau et al., 2015, have also reported the group teaching methods to be superior to individualized teaching methods ^20, 25, 26^. However, the study by Rickheim et al., 2002 depicts no difference in either of the teaching methods ^21^.

High patient satisfaction is associated with efficient communication, personalization of care, patient education, and continuity of care ^27, 28^, whereas a low rating in patient education disrupts the delivery of care and lowers the measure of care outcomes ^29^. As patient satisfaction is mostly subjective, it is measured with the help of the surveys. The patient-satisfaction survey captures self-reported patient evaluations of various points of contact during their medical experience, such as the responsiveness of staff, clinician communication, technical skill, hospital environment, etc. ^30, 31^. Studies by Tung et al., 2009 and Drain, 2001 demonstrates a positive association of patient education and patient satisfaction with the recommendation of a primary care provider to others ^32, 33^. These results are consistent with our study, wherein 92.20% of the participants reported to be very satisfied with the health education activities (post-modification) conducted and would recommend the hospital to others.

As per the pre-implementation data collected, many barriers faced by staff educators in delivering quality health education were highlighted. These include language, place, time of the session, gender, educational material, and presentation skills. A study by Livne et al., 2017 addressed various barriers to patient education experienced by the nurses. The study used the organizational climate theory to examine the potential predictors of barriers to efficient patient education. The study hypothesized that nurses’ perceptions of patient education climate (importance of patient education, based on their daily experience) were related to the barriers of work overload, lack of policies and guidelines, low priority to patient education whereas, the nurses’ role perceptions as patient educators were related to the barriers of difficulty in communicating with patients, insufficient professional knowledge and skills, and the belief that educating patients is not their responsibility. The solutions suggested to reduce barriers included prioritizing patient education, offering a supportive work environment, enabling time for teaching, offering clear guidelines and teaching resources, and developing educating skills for nurses, etc. ^34^. Likewise, the reassessment was effectively modified based on the results of the previous assessment. These included conducting workshops, providing new health education materials, initiating monthly group teaching activities, improvising abilities for health educators, thereby enabling a higher staff satisfaction rate (83.09%). With the modifications, a 100% compliance rate to the educational topics was also achieved compared to a 61.25% compliance rate reported from the previous assessment as a direct result of the unavailability of health education materials.

The PFE activities post-modification had some limitations in terms of inappropriate environment and prolonged or short time of the lecture. Certain similar drawbacks (poor presentation skills, unavailability of educational materials, and poor learning environment) were observed in the previous assessment as well. These are consistent with the limitations faced by Asiri et al., 2013, where a lack of appropriate place and equipment was observed as a barrier. Similarly, DeShazo et al. reported the lack of equipment as a barrier to effective educational intervention ^6, 35^.

The study suggests that through teamwork and maximum utilization of resources, population health management, and patient experience can be enhanced. The study recommends continuous monitoring of patient satisfaction using surveys. Proper coordination among the multidisciplinary team is essential to ensure quality health education.

## Conclusion

Effective health education is a collaborative effort made by the participants and health educators to attain satisfaction. Health educators play an essential role in assisting people to achieve their health goals in a way consistent with their lifestyles, values, and beliefs. Patients and health care providers should be surveyed to assess their experience, satisfaction, and drawbacks associated with them. Thereby facilitating improvisation as and when necessary. By incorporating simple modifications in the educational activities, a higher satisfaction rate can be achieved among the participants and the health educators. Moreover, continuous monitoring and evaluation of health education activities provide effective services to the participants and the community.

## References

1. WHO. Health Education, 2019. Available from: https://www.who.int/healthpromotion/fact-sheet/en/. Accessed on: 06/09/2019.

2. WHO. Health promotion and disease prevention through population-based interventions, including action to address social determinants and health inequity 2019. Available from: http://www.emro.who.int/about-who/public-health-functions/health-promotion-disease-prevention.html. Accessed on: 06/09/2019.

3. WHO. Standards for health promotion in hospitals. Copenhagen 2004. http://www.euro.who.int/_data/assets/pdf_file/0006/99762/e82490.pdf. Accessed on: 06/09/2019.

4. Kemppainen V, Tossavainen K, Turunen H. Nurses’ roles in health promotion practice: an integrative review. Health Promotion International. 2012;28(4):490–501.

5. Schellekens H, Casadevall N. Immunogenicity of recombinant human proteins: causes and consequences. J Neurol. 2004;251 Suppl 2:II4–9.

6. Asiri N, Bawazir AAA, Jradi H. Patients’ satisfaction with health education services at primary health care centers in Riyadh, KSA. Journal of Community Medicine Health Education. 2013;4(1):1–5.

7. JCI. Joint Commission International Accreditation Standards for Hospitals, 2017. Available from: https://www.jointcommissioninternational.org/assets/3/7/JCI_Standards_Only_6th_Ed_Hospital.pdf. Accessed on: 06/09/2019.

8. Oermann MH. Effects of educational intervention in waiting room on patient satisfaction. The Journal of ambulatory care management. 2003;26(2):150–8.

9. Al-Hashem A. Health Education in Saudi Arabia: Historical overview. Sultan Qaboos University Medical Journal. 2016;16(3):e286.

10. Suwaidan H. KFMC Strategic Plan 2015-20202015 23/08/2019. Available from: http://intranet/sites/PSHOC/Stratigic%20Project/KFMC%20Strategic%20Plan%202015-2020%20.pdf#search=KFMC%20Strategic. Accessed on: 06/09/2019.

11. Patient Safety Network Glossary 2019. Available from: https://psnet.ahrq.gov/primers/primer/10. Accessed on: 06/09/2019.

12. Batalden P. Building knowledge for improvement-an introductory guide to the use of FOCUS-PDCA. Quality Resource Group, Hospital Corporation of America Nashville, TN 1992.

13. Bastable SB. Essentials of patient education: Jones & Bartlett Learning; 2016.

14. WHO. Track 2: Health literacy and health behaviour. Health Promotion. 2019. Available from: https://www.who.int/healthpromotion/conferences/7gchp/track2/en/. Accessed on: 06/09/2019.

15. Aghakhani N, Nia HS, Ranjbar H, Rahbar N, Beheshti Z. Nurses’ attitude to patient education barriers in educational hospitals of Urmia University of Medical Sciences. Iranian journal of nursing midwifery research. 2012;17(1):12.

16. ANA. Nursing: Scope and standards of practice: American Nurse Association; 2015.

17. Aghakhani N, Nia, H. S., Ranjbar, H., Rahbar, N., & Beheshti, Z. Nurses’ attitude to patient education barriers in educational hospitals of Urmia University of Medical Sciences. Iranian journal of nursing midwifery research. 2012;17:12.

18. Carneiro SC, Oliveira A, Lopes LJ, Bachion MM, Herdman TH, Moorhead SA, et al. Outpatient Clinic for Health Education: Contribution to Self-Management and Self-Care for People With Heart Failure. International journal of nursing knowledge. 2016;27(1):49–55.

19. Singla DL, Jasser G, Wilson R. Effects of group education on patient satisfaction, knowledge gained, and cost-efficiency in an anticoagulation center. Journal of the American Pharmaceutical Association. 2003;43(2):264–6.

20. Trento M, Passera P, Tomalino M, Bajardi M, Pomero F, Allione A, et al. Group visits improve metabolic control in type 2 diabetes: a 2-year follow-up. Diabetes care. 2001;24(6):995–1000.

21. Rickheim PL, Weaver TW, Flader JL, Kendall DM. Assessment of group versus individual diabetes education: a randomized study. Diabetes care. 2002;25(2):269–74.

22. Best JT, Musgrave B, Pratt K, Hill R, Evans C, Corbitt D. The Impact of Scripted Pain Education on Patient Satisfaction in Outpatient Abdominal Surgery Patients. Journal of PeriAnesthesia Nursing. 2018;33(4):453–60.

23. Abdinnour S. User perceptions towards an ERP system. Journal of Enterprise Information Management. 2015;28(2):243–59.

24. Parnaby P. 2006. [cited 04/09/2019]. Available from: https://www.idea.org/blog/2006/04/01/evaluation-through-surveys/. Accessed on: 06/09/2019.

25. Wilson SR. Individual versus group education: is one better? Patient Education Counseling. 1997;32:S67–S75.

26. Merakou K, Knithaki A, Karageorgos G, Theodoridis D, Barbouni A. Group patient education: effectiveness of a brief intervention in people with type 2 diabetes mellitus in primary health care in Greece: a clinically controlled trial. Health education research. 2015;30(2):223–32.

27. Tevis SE, Kennedy GD, Kent KC. Is there a relationship between patient satisfaction and favorable surgical outcomes? Advances in surgery. 2015;49(1):221.

28. Murdock A, Griffin B. How is patient education linked to patient satisfaction? Nursing2018. 2013;43(6):43–5.

29. Vuong K. Patient education as a quality measure: A Study Based on HCAHPS. 2016.

30. Patient Satisfaction Surveys 2018. Available from: https://catalyst.nejm.org/patient-satisfaction-surveys/. Accessed on: 06/09/2019.

31. Al-Abri R, Al-Balushi A. Patient satisfaction survey as a tool towards quality improvement. Oman medical journal. 2014;29(1):3.

32. Tung Y-C, Chang G-M. Patient satisfaction with and recommendation of a primary care provider: associations of perceived quality and patient education. International Journal for Quality in Health Care. 2009;21(3):206–13.

33. Drain M. Quality improvement in primary care and the importance of patient perceptions. The Journal of ambulatory care management. 2001;24(2):30–46.

34. Livne Y, Peterfreund I, Sheps J. Barriers to patient education and their relationship to nurses’ perceptions of patient education climate. Clinical Nursing Studies. 2017;5(4):65–72.

35. DeShazo JP, Richardson DB, Malotte CK, Rietmeijer CA. Early awareness and uptake of an effective waiting room video intervention by STD clinics. Sexually transmitted diseases. 2011;38(12):1101–6.

